# Constraint-based modeling identifies new putative targets to fight colistin-resistant *A. baumannii* infections

**DOI:** 10.1101/111765

**Authors:** Luana Presta, Emanuele Bosi, Leila Mansouri, Lenie Dijkshoorn, Renato Fani, Marco Fondi

## Abstract

*Acinetobacter baumannii* is a clinical threat to human health, causing major infection outbreaks worldwide. As new drugs against Gram-negative bacteria do not seem to be forthcoming, and due to the microbial capability of acquiring multi-resistance, there is an urgent need for novel therapeutic targets. Here we have derived a list of new potential targets by means of metabolic reconstruction and modelling of *A. baumannii* ATCC 19606. By integrating constraint-based modelling with gene expression data, we simulated microbial growth in normal and stressful conditions (i.e. following antibiotic exposure). This allowed us to describe the metabolic reprogramming that occurs in this bacterium when treated with colistin (the currently adopted last-line treatment) and identify a set of genes that are primary targets for developing new drugs against *A. baumannii*, including colistin-resistant strains. It can be anticipated that the metabolic model presented herein will represent a solid and reliable resource for the future treatment of *A. baumannii* infections.

## Introduction

Bacteria of the genus *Acinetobacter* were long considered harmless, environmental organisms, but from the 1960s onward, an increasing number of reports have documented the emergence of *Acinetobacter* strains of this genus among severely ill, hospitalized patients. These strains showed unusually high levels of resistance to antibiotics that could be used at the time. Also, they gave rise to cross-infections and outbreaks among patients ^1^. Recently, resistance to antibiotics in *A. baumannii* has risen to worrisome proportions (as reviewed in ^2^), from susceptible prior to the 1960s, to multidrug-resistant (MDR) (end 1970s), and extended- and pan-drug resistant (XDR, PDR) today. Currently, *A baumannii* is one of the most prominent organisms that are both antibiotic resistant and involved in health associated infections, the so-called ESKAPE organisms (that include *Enterococcus faecium, Staphylococcus aureus, Klebsiella pneumoniae, Acinetobacter baumannii, Pseudomonas aeruginosa*, and *Enterobacter* spp.^3^).

One of the last-line treatments for MDR *A. baumannii* is colistin, a positively charged molecule that, by interacting with the lipid A moiety of lipopolysaccharide (LPS), causes disorganization of the outer membrane. Unfortunately, colistin resistance in *A. baumannii* has also been reported, thus highlighting the urgency of finding new molecules to face this threat ^4^. Although careful monitoring, antimicrobial stewardship and measures to prevent spread in health care institutions are important for controlling *A. baumannii* infections, new antimicrobial agents and/or strategies are urgently required to eradicate antibiotic resistant strains from affected patients.

To address new possible solutions, a system-level study of antibiotic-response in *A. baumannii* is required. Metabolic network reconstruction and its conversion to a mathematical framework has become a cornerstone for studying the systems biology of metabolism ^5^, allowing the examination of the connection between phenotype and genotype and driving biological discoveries. In particular, constraint-based tools (such as Flux Balance Analysis, FBA) enable the estimation of the rate that metabolites’ flow through a metabolic network and to compute cellular phenotypes for various growth conditions ^6^. Interestingly, by identifying those genes whose deletion is predicted to impair cellular growth, this *in silico* technique can be used to predict essential genes (EGs) at a genome-scale. Following a metabolic modelling approach, several EGs datasets have already been derived for important pathogens such as *Helicobacter pylori* ^7^, *Pseudomonas aeruginosa* ^8^, *Mycobacterium tuberculosis* ^9^, and *Staphylococcus aureus* ^10^. Usually, such predictions are performed simulating growth in an arbitrarily defined medium, accounting for the main nutrients used by the microbe and without imposing any additional constraint to the model. Indeed, the search space of essential genes predicted can be narrowed by imposing additional constraints on the model. One possibility consists of modulating the flux admissible across each reaction on the basis of the expression values of the corresponding genes. By doing so, it is possible to generate context-specific models that reflect the actual set of reactions employed ^11^. This approach promises to reduce i) the gap between the predicted and real cellular metabolic landscapes, and ii) the number of false positives/negatives in EGs predictions. Additionally, it might reveal hints for the synergistic use of antibiotics and, in particular, to the possible additional targets that might arise from the adaptation/response of a microbe’s metabolism to a single antibiotic. Indeed, changes in gene expression might redirect the cellular metabolic fluxes in such a way that novel and untapped essential reactions may emerge, representing good candidates for a synergic antibiotic. Despite that the use of antibiotics in combination is sometimes questionable, this approach can be considered in cases of severe infections and it has been shown to be effective in the case of *Pseudomonas* and *Acinetobacter* spp.^12 13^.

Here, we explored the system-level metabolic consequences of *A. baumannii* exposure to colistin. We integrated gene expression data during exposure to colistin ^14^ with a newly reconstructed genome scale metabolic model, allowing for constraint-based modelling of the type strain ATCC 19606. Our data revealed the metabolic reprogramming that occurred in this strain following the establishment of a stressful condition such as the presence of an antibiotic. Furthermore, the metabolic reconstruction provided here represents an important resource for the future understanding of *A. baumannii* metabolism and for the detection and identification of novel drug targets.

## Results and Discussion

### Genome-scale *A. baumannii* ATCC 19606 model is consistent with large scale phenotypic data

A preliminary draft reconstruction of the *A. baumannii* ATCC 19606 metabolic model was obtained through the Kbase server (http://kbase.us). This was manually curated as described in Methods. Afterwards, we used previously published large-scale phenotypic data^15^ to validate our reconstruction over a large set of experimental tests. Manual curation was performed by comparing FBA outcomes with such auxotrophies data (determined through Phenotype Microarray (PM) technology). During this process, the capability of our model to represent the observed phenotypes was tested.

Growth rates were firstly estimated *in silico* in simulated Simmons minimal medium (a standard bacteriological medium that contains only essential inorganic salts) under aerobic conditions by iteratively probing each C-source used in PM plates. During these simulations, biomass optimization was selected as the model objective function (O.F.). Results of the simulations (either “growth” or “no growth”, *i.e*. the estimated flux value across biomass assembly reaction) were compared with the activity directly measured during an experimental phenotype microarray experiment, and discrepancies identified between the *in silico* and experimental data were manually adjusted as possible (such as by filling in missing transport reactions or metabolic gaps).

Following this procedure, we reached an overall agreement of about 88% between the *in silico* and experimental data: out of the 67 *in silico* screened metabolites, 25 were correctly found to be carbon and energy sources for *A. baumannii* ATCC 19606 (true positives) and 34 not (true negatives) while only 8 disagreements remained, 3 false negatives and 5 false positives. All data are briefly summarized in Figure 1, and a detailed description of the outcomes of the comparison is reported in Supplementary Material S1, Supplementary Table 1.

**Figure 1:**
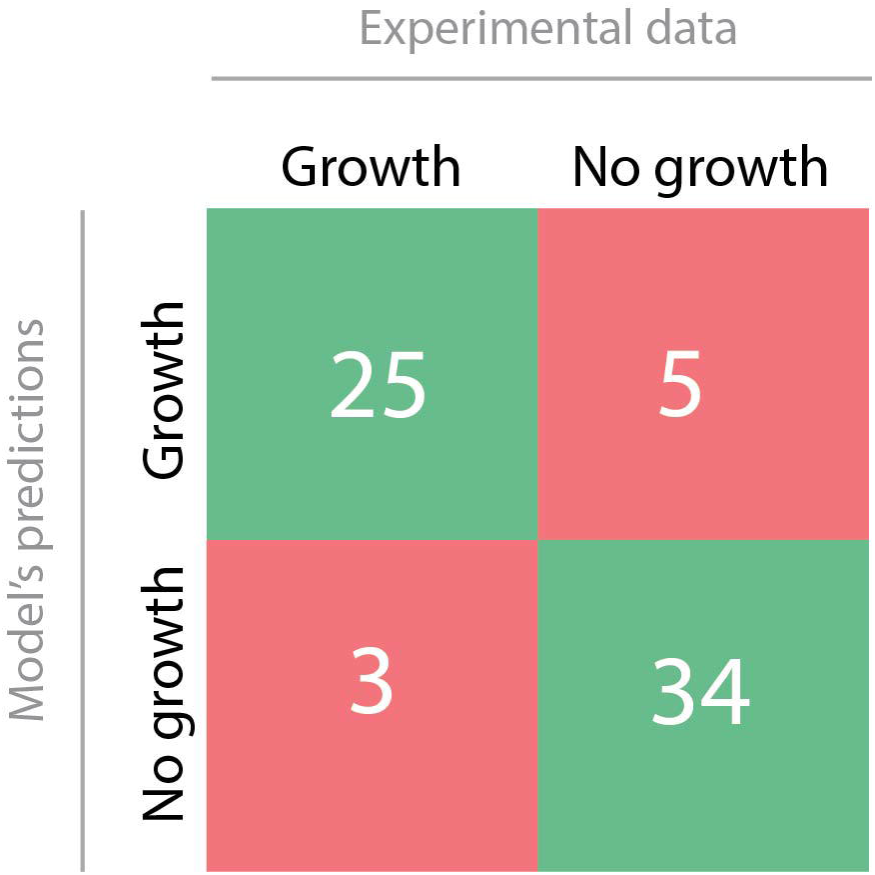
Comparison between *in silico* and wet-lab experimental outcomes.

The current version of the *A. baumannii* ATCC 19606 genome-scale metabolic model (named iLP844 according to the current naming convention ^16^) contains 1615 reactions (162 exchange reactions), 1509 metabolites, and 844 genes (~23% of all ORFs present in this organism, see Supplementary Material S2). Importantly, this proportion is comparable with the coverage of *Escherichia coli* K12 model, iAF1260 ^17^ (27%), *A. baylyi* ADP1 model, iAbaylyi ^18^ (22%), and *A. baumanni* AYE model, AbyMBEL891 ^19^ (17%). A COG classification of all the genes embedded in the model is provided in Table 1. The final *A. baumannii* ATCC 19606 model is available as supplementary material in both validated SBML and JSON formats, the latter embedding cross-references to several databases (Supplementary Material S3 and S4, respectively).

**Table 1:** Number of genes in the model per COG categories.

| COG<br>Functional<br>Category | Description | N. of<br>genes |
| --- | --- | --- |
| J | Translation, ribosomal structure and biogenesis | 28 |
| A | RNA processing and modification | 0 |
| K | Transcription | 11 |
| L | Replication, recombination and repair | 21 |
| B | Chromatin structure and dynamics | 0 |
| D | Cell cycle control, cell division, chromosome partitioning | 1 |
| Y | Nuclear structure | 0 |
| V | Defense mechanisms | 4 |
| T | Signal transduction mechanisms | 5 |
| M | Cell wall/membrane/envelope biogenesis | 60 |
| N | Cell motility | 0 |
| Z | Cytoskeleton | 0 |
| W | Extracellular structures | 0 |
| U | Intracellular trafficking, secretion, and vesicular transport | 0 |
| O | Posttranslational modification, protein turnover, chaperones | 20 |
| C | Energy production and conversion | 120 |
| G | Carbohydrate transport and metabolism | 61 |
| E | Amino acid transport and metabolism | 182 |
| F | Nucleotide transport and metabolism | 57 |
| H | Coenzyme transport and metabolism | 82 |
| I | Lipid transport and metabolism | 97 |
| P | Inorganic ion transport and metabolism | 71 |
| Q | Secondary metabolites biosynthesis, transport and catabolism | 16 |
| R | General function prediction only | 32 |
| S | Function unknown | 12 |
| X | No Functional Class Found | 15 |

### Constraint-based modelling identifies metabolic EGs

As already mentioned, the identification of EGs is one of the key-steps in a drug discovery pipeline. Indeed, both general and condition-specific EGs can be distinguished ^20^. The formers are required to sustain life under virtually all growth conditions; conversely, under specific constraints, changes of central metabolism may occur leading not only to a change in flux distribution throughout the network, but may also lead to changes in gene essentiality and the appearance of condition-specific essential genes. Hence, we systematically evaluated relevant switches in both unconstrained and constrained models (different *scenarios*), by imposing the necessary constraints to the metabolic reconstruction. Accordingly, screens for EGs were performed for multiple specific conditions: starting by simply changing the set of available nutrients (i.e. simulating different environmental niches) and then by simulating stressful situations such as antibiotic exposure and mutations (using available expression data in such conditions).

#### Nutrient availability influences identified metabolic EGs

As we were interested in modelling the system in a gradually constrained manner, we initially simulated an arbitrary rich medium, allowing our model to have virtually all the nutrients needed - as likely happens inside of a host ^19^. To do so, we set the lower bound of exchange reactions as described in methods. Then we performed *in silico* gene deletions for each gene in the model. Accordingly, each gene was defined as essential if its elimination destroyed the network’s ability to synthesize at least one key biomass molecules (*i.e*. the model predicts no-growth); otherwise, the gene was considered to be dispensable. Gene essentiality analysis was performed through both FBA and MOMA approaches (see Methods), which both lead to the identification of 57 EGs. Figure 2A shows the values of the ratio between the predicted growth rate of the gene knock-out mutant and the wild type strain (GR ratio, see Methods). The complete list of EGs and their functions is reported in Supplementary Material S5.

Next, we repeated EGs prediction by simulating growth in a minimal medium (Simmons medium, as defined in Supplementary Material S1, Supplementary Table 2). As shown in Figure 2B, this approach labelled a total of 125 genes as indispensable for growth in this condition (see Supplementary Material S5 for the complete list). Differences emerging from these two simulations highlight how nutrient availability affects cell metabolism and, interestingly, how different environmental pressures influence gene essentiality. Particularly, 57 genes were predicted to be essential under both the tested growth conditions, while 68 are likely to become essential only when limited nutrient availability force the cell to reprogram its metabolic behaviour (*i.e*. in Simmons medium, Figure 3A).

**Figure 2:**
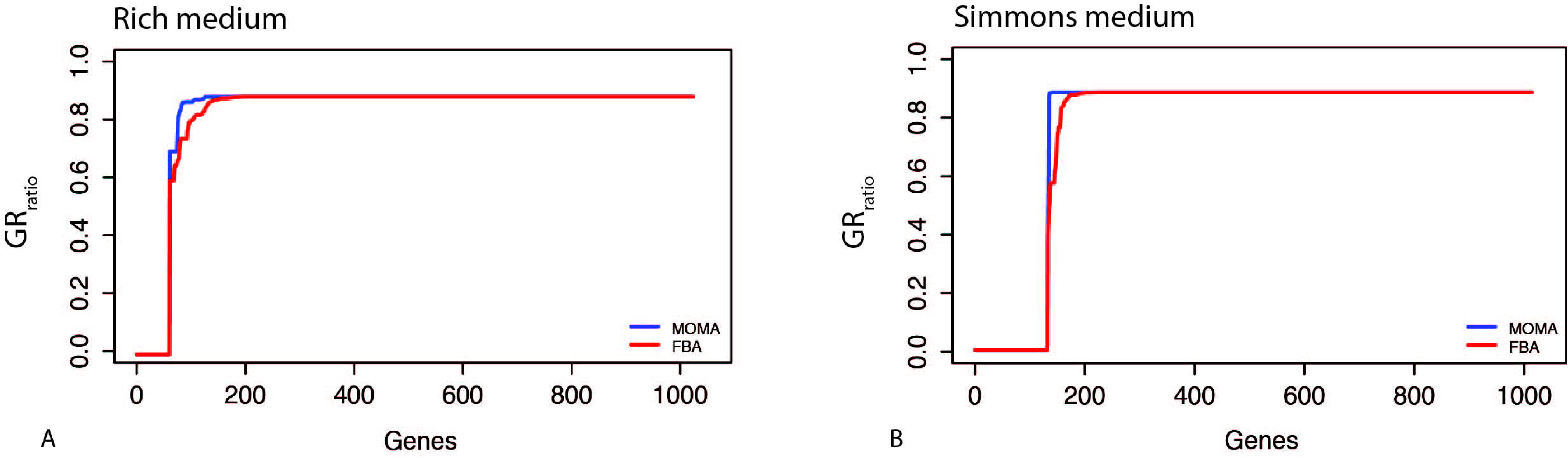
GR_ratio_ value for each gene deletion in rich (A) and minimal (B) media. Blue and red lines represent MOMA and FBA predictions, respectively. Please note that, in order to make the analysis more comprehensive, also gap-filling genes (i.e. those virtually coding for gap-filling reactions) were included, leading to a total of 1043 simulated knock-outs.

**Figure 3:**
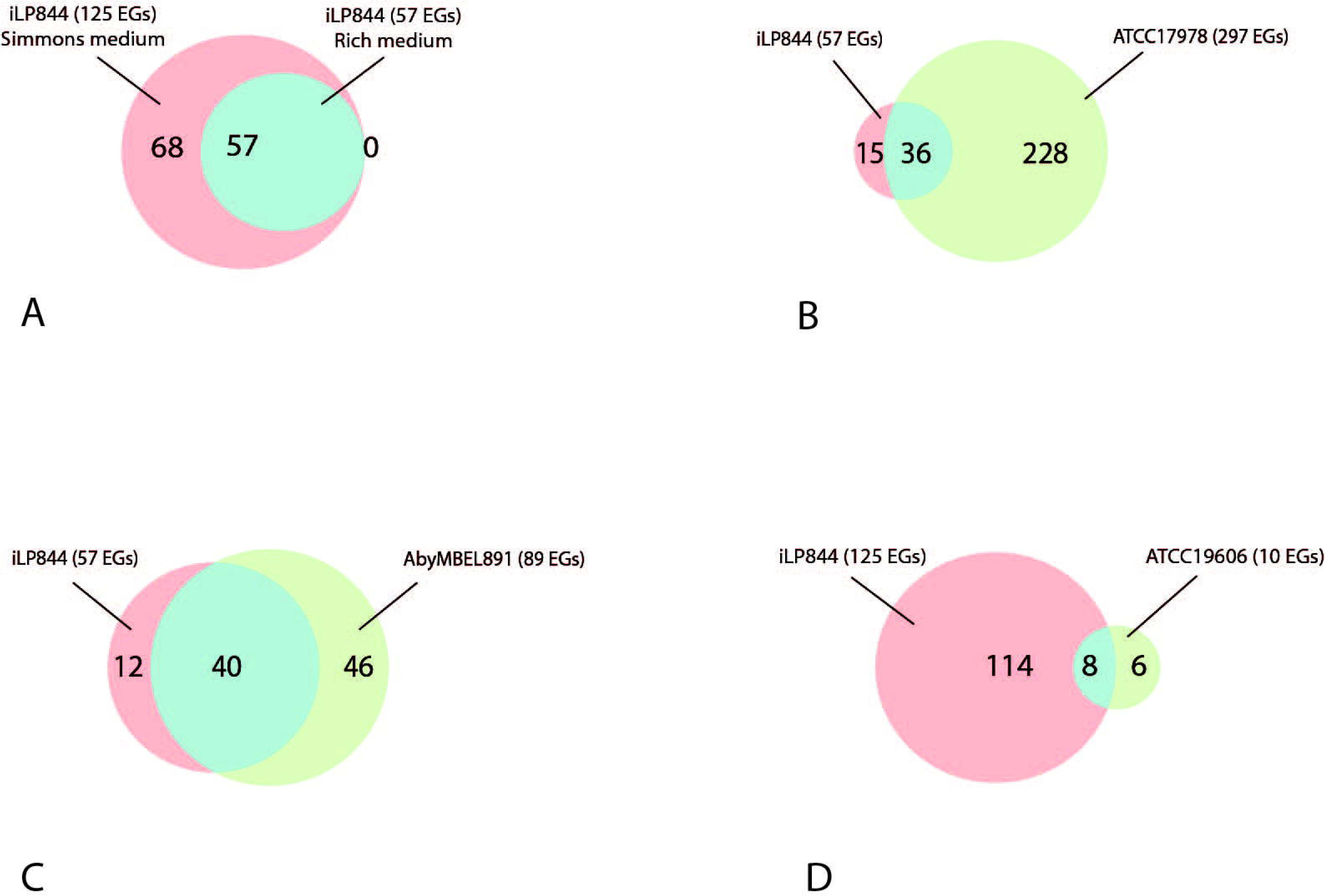
A) A venn diagram proportionally showing EGs predicted only in Simmons medium (pink), EGs predicted only in rich medium (green), and EGs predicted by both (blue). B) A venn diagram proportionally showing EGs predicted *in silico* only by iLP844 (pink), EGs obtained only by wet-lab experiment in ATCC 17978 (green), and EGs predicted by both methods (blue). C) A venn diagram proportionally showing essential reactions predicted in iLP844 (pink), essential reactions predicted only in *A. baumannii* AYE model (AbyMBEL891) (green), and essential reactions predicted by both (blue). D) A venn diagram proportionally showing EGs predicted only in iLP844 (pink), EGs predicted only by wet-lab experiment in *A. baumannii* ATCC 19606 cell (green), and EGs predicted by both (blue).

#### Predicted EGs are consistent with available experimental datasets

A large body of data exists concerning *A. baumannii* gene essentiality. Here we used such information both to validate our EGs prediction and to understand whether the identified EGs sets are particular to *A. baumannii* ATCC 19606.

First we compared the EGs dataset obtained in the arbitrary rich medium to that obtained with an *in vivo* experiment on *A. baumannii* ATCC 17978 pathogenesis ^21^. By using LB medium (a well-known bacteriological rich medium), Wang et al.^21^ labelled 481 genes as essentials for that strain. However, not all of them were comparable with our predictions since a large fraction was neither metabolic nor possess an orthologous gene in *A. buamannii* ATCC 19606. In both cases these genes are absent in iLP844. For the same reason, not all the 57 EGs found through our simulation were comparable with the reported experiment. After performing all these necessary restrictions, we reached the result shown in Figure 3B, *i.e*. 36 genes have been predicted to be essential by both approaches (*in silico* and wet-lab) for the two *A. baumannii* strains considered (Figure 3B). A complete description of these EGs is provided in Supplementary Material S5 and represents an experimentally validated dataset in the context of *A. baumannii* drug target identification. Nevertheless, the two experiments show large discrepancies. The most likely reason for such inconsistency is strains genomic diversity, as previously reported for *E. coli* strains ^22^.

Furthermore, our predictions in arbitrary rich medium were compared to those achieved performing the same analysis on the AbyMBEL891 model, an existing model of *A. baumannii* AYE^19^. In order to implement the simulation, it was necessary to perform a preliminary editing step on the AbyMBEL891 model, as the entire set of gene-reaction-rules was missing from the main reconstruction file. This difficulty in running the analysis highlights the need for a common protocol to be adopted during metabolic reconstruction and a standard to be reached in order to facilitate model re-use and data sharing among research groups. Nevertheless, after including the genes in the Abymbel891 model, we carried out single gene deletion analysis on both models, as described in the methods. As shown in Figure 3C, 40 genes were predicted to be essential in both models, whereas 12 and 46 EGs were specific for *A. baumannii* ATCC 19606 and AYE, respectively. Information about the gene function are reported in Supplementary Material S5.

Comparisons were also carried out between our *in silico* predictions and wet-lab results in minimal (Simmons) medium. Specifically, we compared our EGs set to that obtained by Dorsey et al. through insertional mutagenesis experiments with *A. baumannii* ATCC 19606^23^, where the metabolic deficiency of insertion derivatives was subsequently confirmed, identifying essentiality of 10 disrupted genes. Repeating the assay *in silico*, our model correctly represented the phenotypes of the *A. baumannii* mutants, with 8 out of the 10 genes predicted as essential by Dorsey and colleagues also shown to be essential in iLP844 (Figure 3D, Supplementary Material S5). Additionally, in 6 out of the 8 cases, *A. baumannii* ATCC 19606 model growth was correctly restored (as done in the corresponding wet-lab experiments) by adding to the minimal medium the metabolite(s) whose production was affected by the mutation.

### Antibiotic treatment defines condition-specific models

Although a large fraction of the predictions was supported by previous experimental data, a possible source of error, using the methodology described above, stems from the observation that not all the reactions of the model will be active during growth in a given physiological condition. In particular, changes in gene expression are likely to influence the activity rate of the corresponding cellular metabolic reactions, leading to the observation that a given reaction can be considered ‘turned on’ or ‘off on the basis of the expression levels of the encoding gene(s). Using available computational methodologies, it is possible to modulate the flux across each reaction on the basis of the expression values of the corresponding genes. This allows for taking a picture of the current metabolic state and tightening up the predictive capabilities of the model itself. Accordingly, as the dynamic changes of metabolic reprogramming are likely mirrored by changes in gene essentialities, a possible solution for avoiding or reducing false positives is to merge transcriptomics data of the tested *scenario* into the genome-scale model.

Arguably, one of the most interesting physiological conditions of *A. baumannii* strains is the exposure to antibiotics and to colistin in particular^14^. Importantly, both the (metabolic) consequences and the occurrence of targets to be used in a synergic treatment are, currently, almost untapped. In order to study the dynamic changes of the metabolic network following antibiotic exposure and to derive a more realistic picture of gene essentiality patterns in a real scenario (antibiotic treatment), we used available transcriptomic data for *A. baumannii* ATCC 19606 in response to colistin treatment ^14^. Up-regulation and down-regulation ratios (and corresponding *P*-values) of genes were combined with the iLP844 by using MADE (Metabolic Adjustment by Differential Expression)^24^. Briefly, MADE uses statistically significant changes in gene expression measurements to determine binary expression states (highly and lowly expressed reactions) *i.e*. reactions are turned on and off depending on the changes in mRNA transcript levels. Thus, by mapping gene expression data into the model, the *in silico* metabolic predictions are more consistent with the actual physiological state of the cell.

In the experiment by Henry et al. ^14^, *A. baumannii* was grown in two different media, *i.e*. with and without 2 mg/L of colistin, and then sampled at 15 and 60 minutes after exposure. Following the described approach, we integrated the available transcriptomic data regarding all the metabolic genes embedded in our *in-silico* reconstruction (*i.e*. about 80 genes). Accordingly, we obtained four distinct models, each representing the predicted functional metabolic state of the cell at both 15 and 60 minutes, treated and untreated with colistin. These models differ in that some of their reactions are (completely) ‘turned on’ or ‘off’ according to the measured levels of their corresponding genes. Afterwards, optimization of the four models was performed, allowing the analysis of flux distribution in the network and the occurring metabolic reshape.

### Colistin exposure changes predicted metabolic fluxes in central *A*. *baumannii* pathways

In order to highlight changes in the overall metabolic behaviour and to identify changes on the metabolic rewiring occurring after antibiotic exposure, we compared flux distributions at the two time-points by calculating the flux ratio (RF_ratio_, see Methods) of treated *vs*. untreated models, for all the reactions. However, FBA only provides one of the possible optimal solution out of many alternative (and feasible) cellular flux distributions. Hence, in order to correctly predict metabolic changes following antibiotic exposure we restricted the feasible solution space by performing Flux Variability Analysis (FVA) ^25^. This approach allows estimating the minimum and maximum flux admissible across each reaction (under the same constraints as in FBA) and hence it can be used to estimate the correctness and accuracy of FBA predictions (see Methods).

As shown in Supplementary Material S6, according to FVA, the range of admissible flux is sometimes very large (spanning from the minimum −1000 to maximum 1000 mmol/g*h^−1^ in some cases), revealing the lack of accuracy in some of FBA-derived predictions. We here used FVA outcomes (as described in Methods) to filter out those reactions whose fluxes display little variation. In other words, each reaction was considered for downstream analyses only if both the maximum and minimum FVA predicted fluxes did not differ from the FBA predicted flux by more than 20%. Consequently, we were left with 901 reactions at 15 minutes and 970 reactions at 60 min. It is worth noting that several intervals of admissible flux ranges were tested and we report in Supplementary Materials S1, Supplementary Figure 1 the number of reactions filtered for each set of intervals. After carrying out this preliminary step, we observed the effects of the treatment at the metabolic level (for each reaction) by comparing the flux values in the untreated *vs*. treated condition.

Both qualitative and quantitative flux changes were analysed by dividing the reactions into three categories (‘steady’, ‘increasing’, and ‘decreasing’, see Figure 4) according to their trends in the examined experimental conditions. Also, we report a survey of the pathways in which they are involved in and their relative abundance for each category. Reactions’ fluxes were considered ‘steady’ if their values did not change in the two conditions, otherwise they were defined to be ‘increasing’ or ‘decreasing’ according to the corresponding trend.

As shown in Figure 4, at both 15 and 60 min time-points there is an increase in flux in most of the reactions. Interestingly, such change in flux mainly occurs in three biosynthetic pathways: fatty acid, peptidoglycan, and lysine biosynthesis. On the other hand, under the given constraints, there is a change in flux in some catabolic pathways (mainly involved in sugars and nucleotide metabolism). In our opinion, such a finding could be related to the rearrangement of the external membrane layer, a well-known effect of colistin treatment. If this is true, it is possible that the cell reacts to the antibiotic treatment by trying to repair the damage established by colistin while at the same time redirecting a certain amount of LPS components to catabolic processes.

**Figure 4:**
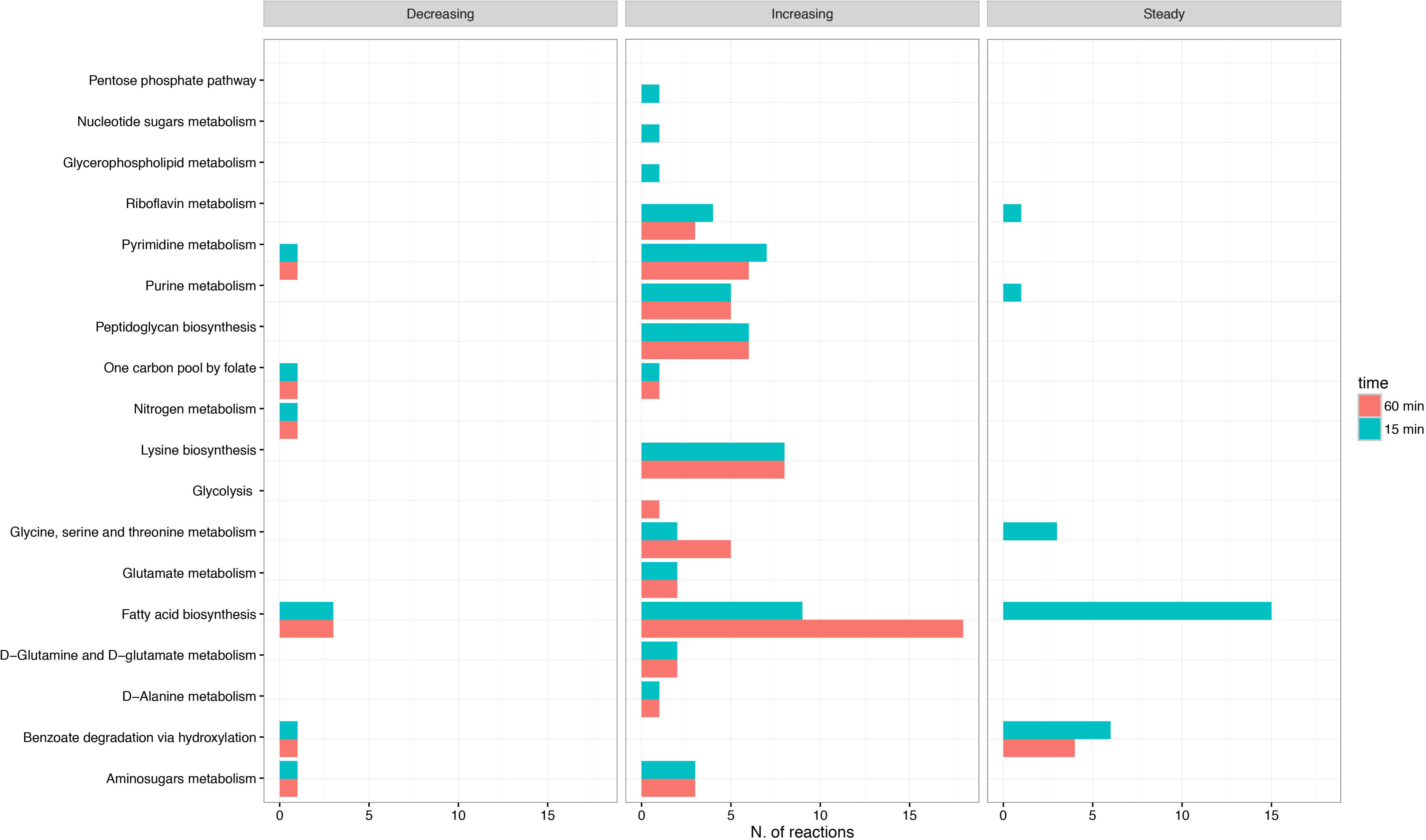
Abundance plot of reactions affected by colistin treatment at 15 (blue) and 60 (pink) minutes arranged according to three categories: ‘steady’, ‘increasing’, ‘decreasing’. Pathways which they belong to are reported.

Also, we would like to point-out that, although a down-regulation of certain genes involved in fatty acids biosynthesis was detected by Henry et al., here our data suggest that it does not necessarily imply a turning-off of the pathway. On the contrary, in our simulation fatty acid biosynthesis registers an increase in flux, probably as a side-effect of LPS disassembly as stated above.

### Colistin exposure changes gene essentiality patterns

According to the new constraints taken into account, gene essentiality was re-evaluated by calculating growth ratios (FBA and MOMA) at both 15 and 60 minutes after exposure to colistin. As for the analysis involving nutrient availability, shifts in gene essentiality emerged following antibiotic stress. The complete sets of the predicted EGs for each condition have been reported in Supplementary Material S5. As with the previous case, we can easily recognize genes likely to be essential in both conditions (treated and not) and, more interestingly, genes that emerged as essential only after the treatment. Specifically, following 15 minutes of colistin exposure, a total of 65 EGs were predicted: 57 were required both in presence and absence of colistin, but an additional 16 EGs were marked as condition-specific: 8 related to the non-treated model and 8 related to the treated one, reported in Table 2 and in Figure 5A.

**Figure 5:**
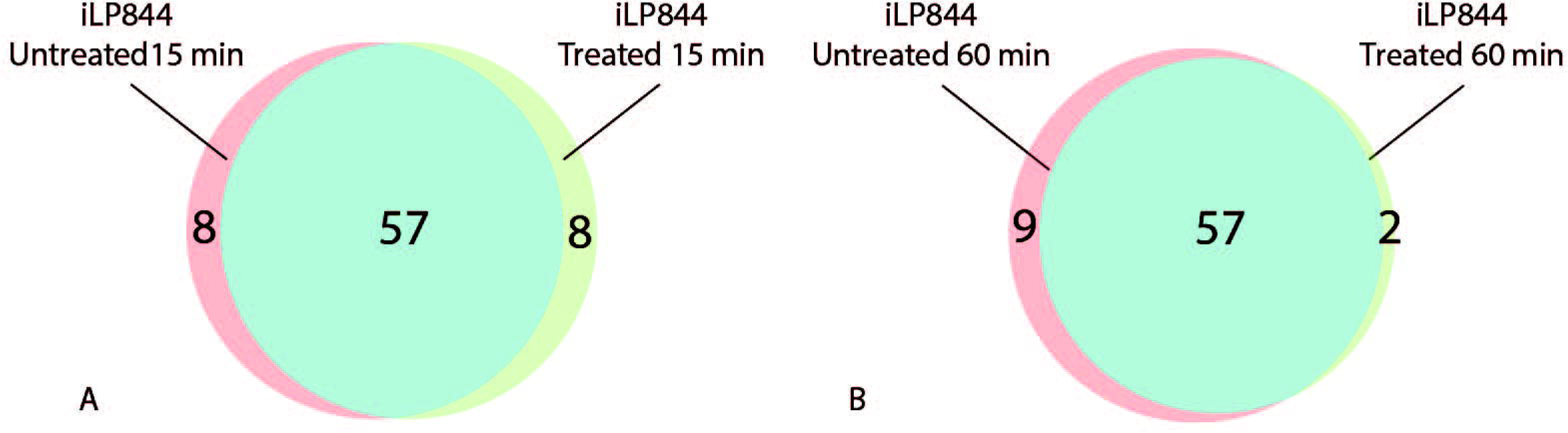
Venn diagrams proportionally showing EGs predicted only in absence of colistin (pink), EGs predicted only in presence of colistin (green), and EGs predicted by both (blue), at 15 minutes (A) and 60 minutes (B).

**Table 2:** Complete set of condition specific EGs found.

| Locus Tag | Function | 15'<br>without<br>colistin | 15'<br>with<br>colistin | 60'<br>without<br>colistin | 60'<br>with<br>colistin | LPS-without<br>colistin | LPS-with<br>colistin |
| --- | --- | --- | --- | --- | --- | --- | --- |
| HMPREF0010_00435 | phosphopyruvate hydratase | Yes | No | Yes | No | No | No |
| HMPREF0010_00813 | fructose-bisphosphate<br>aldolase, class II, Calvin<br>cycle subtype | Yes | No | Yes | No | No | No |
| HMPREF0010_00815 | phosphoglycerate kinase | Yes | No | Yes | No | No | No |
| HMPREF0010_00975 | amino acid ABC<br>transporter periplasmic<br>protein | Yes | No | Yes | No | No | No |
| HMPREF0010_01733 | PAP2 superfamily protein | Yes | No | Yes | No | No | No |
| HMPREF0010_01995 | 2,3-bisphosphoglycerate-<br>independent<br>phosphoglycerate mutase | Yes | No | Yes | No | No | No |
| HMPREF0010_02140 | fructose-1,6-bisphosphatase | Yes | No | Yes | No | No | No |
| HMPREF0010_03273 | glucose-6-phosphate<br>isomerase | Yes | No | Yes | No | No | No |
| HMPREF0010_00342 | ornithine<br>carbamoyltransferase | No | Yes | No | No | No | Yes |
| HMPREF0010_00949 | malonate decarboxylase,<br>epsilon subunit | No | Yes | No | Yes | No | No |
| HMPREF0010_01149 | acetyl-CoA carboxylase,<br>biotin carboxylase | No | Yes | No | Yes | No | No |
| HMPREF0010_01969 | argininosuccinate lyase | No | Yes | No | No | No | Yes |
| HMPREF0010_02047 | carbamoyl-phosphate<br>synthase, large subunit | No | Yes | No | No | No | Yes |
| HMPREF0010_02048 | carbamoyl-phosphate<br>synthase, small subunit | No | Yes | No | No | No | Yes |
| HMPREF0010_02972 | argininosuccinate synthase | No | Yes | No | No | No | Yes |
| HMPREF0010_03445 | 3,4-dihydroxy-2-butanone<br>4-phosphate synthase | No | Yes | No | No | No | No |
| HMPREF0010_02330 | glutamine synthetase, type<br>I | No | No | Yes | No | No | No |
| HMPREF0010_00434 | 3-deoxy-8-phosphooctulonate<br>synthase | No | No | No | No | Yes | No |
| HMPREF0010_03552 | phenylphosphate<br>carboxylase, delta subunit | No | No | No | No | Yes | No |
| HMPREF0010_03553 | sugar phosphate isomerase | No | No | No | No | Yes | No |
| HMPREF0010_00392 | ornithine-oxo-acid<br>transaminase | No | No | No | No | No | Yes |
| HMPREF0010_00419 | N-acetyl-gamma-glutamyl-phosphate reductase | No | No | No | No | No | Yes |
| HMPREF0010_01215 | ArgJ protein | No | No | No | No | No | Yes |
| HMPREF0010_01331 | ribose-phosphate<br>pyrophosphokinase | No | No | No | No | No | Yes |
| HMPREF0010_01382 | acetylglutamate kinase | No | No | No | No | No | Yes |
| HMPREF0010_01861 | glutamate racemase | No | No | No | No | No | Yes |

The same outline has been depicted in the second time-point condition (60 minutes): we identified 9 and 2 condition-specific EGs in the absence and in the presence of the antibiotic, respectively, see Table 2. Moreover, we found the same set of 57 EGs mentioned above (see Figure 5B), suggesting that this represents a functionally relevant set of genes for sustaining growth in *A. baumannii* ATCC 19606. Interestingly, however, some genes switch from the ‘essential’ condition to the ‘nonessential’ one, following the exposure to colistin.

The two new sets of EGs show how changes in gene expression induced by the presence of the antibiotic might influence gene essentiality patterns in the strain ATCC 19606 and provide additional, nontrivial targets for drug design in such organism. We also performed additional robustness analyses in order to test whether nutrients depletion occurring in treated and untreated samples during the *in vivo* experiments could impact the set(s) of predicted EGs. Specifically, the robustness of the number of predicted EGs in each of these conditions (*i.e*. treated *vs*. untreated samples) in respect to possible variations in the medium composition was assessed via random permutation. We tested up to 1,000 different nutritional compositions as described in detail in Supplementary Material S1, Supplementary Figure 2). The results showed that possible changes in the nutritional environment had only minor implications for the set of predicted EGs.

Further, with the aim of discriminating whether the products of all the identified hypothetical EGs are *A. baumannii* specific or have orthologs in *Homo sapiens*, meaning they would not represent good candidates for antibiotic treatment development, the sequences of the identified potential EGs were used to probe the human genome. Based on this BLAST ^26^ search (see Methods), we excluded from further studies those genes presenting more than 30% sequence identity with their human counterparts. Targeting of such genes is non-ideal since they may cause potential side-effects by perturbing critical components in the human body. All BLAST results are reported in Supplementary Material S7.

Among all the queries, we identified 40 (out of 57) general EGs and 3 (out of 8) condition-specific EGs that do not have any human orthologous. Thus, the 40 EGs represent valuable targets for further development of brand new drugs against *A. baumannii* ATCC 19606 infections. However, it is relevant to remark that, while these 40 general EGs could have been detected in several conditions, the other 3 condition-specific EGs are the result of specific constraints integrated in the model (gene expressions data). Hence, as already mentioned, they are nontrivial detections and they could represent a suitable horizon in the field of colistin-coupled treatment. The three genes, named HMPREF0010_00949, HMPREF0010_02972 and HMPREF0010_03445, respectively encode a malonate decarboxylase (epsilon subunit), an arginine succinate synthase and a 3-4-dihydroxy-2-butanone-4-phopsphate synthase. Interestingly, malonate decarboxylase epsilon subunit has already been characterized in the closely related organism *Pseudomonas putida* and labelled as an indispensable component of the enzyme for the cyclic decarboxylation of malonate^27^. However, to the best of our knowledge, no therapies targeting this protein have been developed to date. The product of 3-4-dihydroxy-2-butanone-4-phopsphate synthase is an intermediate in the biosynthesis of riboflavin. The enzyme requires a divalent cation, preferably Mg_2_^+^, to be active. The step becomes essential after colistin treatment as the antibiotic is predicted to cause an increase in flux through this pathway, probably following the shutdown of other parts of the network due to the down regulation of the corresponding genes. The last enzyme, the arginine succinate synthase, is an enzyme catalysing the penultimate step in arginine biosynthesis (urea-cycle): the ATP-dependent ligation of citrulline to aspartate in order to form arginino-succinate, AMP, and pyrophosphate.

### EGs in colistin resistant *A. baumannii*

Up to now, we have presented how metabolic reconstruction and mathematical modelling can be used to explore the strain’s metabolic response during colistin treatment and how it can lead to the identification of novel potential drug targets. Our last attempt is now to illustrate how, starting from the same available experimental data, the model can be employed as a ready-to use blueprint in order to test new hypothesis.

As it was reported by Moffatt et al. ^28^, the mechanism responsible for colistin resistance is linked to LPS. Specifically, mutations in the *lpxA, lpxC*, and *lpxD* genes have been reported as the main cause of LPS loss, thus abolishing the initial charge-based interaction with the antibiotic. Hence, to simulate an *A. baumannii* LPS (LPS^−^) deficient and colistin-resistant strain, we removed this component from the biomass formulation in our genome-scale model. Then, to determine which genes are central for the cell’s survival in such a condition, we used the transcriptomic data of the mutant strain in the presence/absence of colistin at 60 minutes ^14^ and mapped the data onto the new LPS^-^ model. After this, we repeated the EGs prediction pipeline described above.

The analysis yielded a total of 62 and 70 EGs in the untreated and treated condition, respectively. Even in this case, the two sets share some elements (59 EGs) that remain mandatory for the cell in the two conditions (listed in Supplementary Material S5). Additionally, it is possible to observe that 11 genes become essential (reported in Table 2) only after antibiotic exposure: 5 of them were already found to be EGs in the wild type strain while 6 represent specific EGs of the mutant. Since the latter are non-trivial EGs (obtained only through gene expression integration into the model) they have been re-used as seed for an additional BLAST search against the human genome (see Supplementary Material S7). The search led to the identification of 4 genes that do not have orthologs in humans: HMPREF0010_01215 encoding for glutamate-N-acetyltransferase (member of the ornithine acetyltransferase, OAT, family), HMPREF0010_00419, encoding for N-acetyl-gamma-glutamyl-phosphate reductase, HMPREF0010_02972 (previously described), and HMPREF0010_01382 encoding for N-acetyl-L-glutamate-kinase, all of which are involved in the arginine biosynthesis pathway, and HMPREF0010_01861 that encodes for a glutamate-racemase. This group of 4 EGs represents a potential achievement obtained from this work as it suggests specific targets to be taken into consideration when developing therapies in combination with colistin.

#### Predicted EGs are common in *A. baumannii*

Finally, we checked the distribution of EGs predicted for the strain ATCC 19606 within the entire *A. baumannii* species. The sets of predicted EGs were searched in all of the 1099 *A. baumannii* genomes sequenced to date, as described in the Methods. The overall result is shown in Supplementary Material S1, Supplementary Figure 3. The general trend observed was that more than 90% of the genomes analysed possessed the searched queries (identity >30%). Also, our analysis shows that this tendency is kept almost unchanged even when imposing an identity threshold greater than 50%, 70% and 90%. Accordingly, it can be stated that the possible target genes are broadly distributed and their sequence is conserved at the *A. baumannii* species level. Although we do not have any information about the EGs at such a wide level, this preliminary result is encouraging, since it expresses the possibility that the target genes we indicated for *A. baumannii* ATCC 19606 are probably common targets in most of *A. baumannii* type infections.

## Conclusions

In this work, we have reconstructed and validated a genome-scale metabolic model of *A. baumannii* ATCC 19606. The model is comprehensive and accurate, as it covers ~23% of all CDSs in the genome of this microorganism and it was shown to have 88% agreement with Phenotype Microarray growth experiments. Based on the model’s reliability, we applied constraint-based modelling to derive a global understanding of the behaviour of this metabolic system. By integrating gene expression data with constraint-based modelling we described the metabolic reprogramming occurring after colistin-exposure in *A. baumannii* and the changes in the pattern of gene essentiality during this stress condition. All the sets of condition-specific putative target genes that we propose have been compared (and partially validated) with the results obtained from experiments found in the literature. Some of these genes, although not yet experimentally validated, might represent primary targets for future research on the treatment of both the wild type and LPS-mutant (*i.e*. colistin resistant) strains. Our results have practical implications for the identification of new therapeutics as the identified essential genes can be used in drug-design pipelines. Moreover, we showed that the sequences of predicted EGs for the type strain ATCC 19606 are shared by most of the members of *A. baumannii* species, encouraging further research to check whether they are valuable drug targets for a larger number of strains than currently known. Finally, it can be anticipated that the iLP844 model illustrated herein represents a reliable and solid platform for further developments and the system-level understanding of the physiology of *A. baumannii* representatives and for the treatment of their infections.

## Methods

### Draft model reconstruction

We obtained a draft metabolic model of *A. baumannii* ATCC 19606 based on the genome annotation using Kbase automated reconstruction method (https://kbase.us/) ^29^. This reconstruction was then thoroughly inspected following the main steps listed in Thiele and Palsson ^5^, and refined by integrating data from additional functional databases (MetaNetX, Bigg, Seed, KEGG). Further integration was performed by searching for orthologous genes (genes likely having an identical biological function in a different organism) in closely related organisms (*Acinetobacter baumanii* AYE, *Acinetobacter baylyi* ADP1, and *Escherichia coli*) through a BBH (Bidirectional Best Hit) approach (inParanoid^30^). Information regarding transport proteins was obtained probing the Transporter Classification Data Base (TCDB ^31^) and transportDB ^32^.

In order to predict proper phenotypes, the general biomass producing reaction of Gram negative bacteria automatically generated by Kbase was substituted with a more accurate one that takes into account strain’s specific components, which was recovered from the previously reported model of the related strain *A. baumannii* AYE, AbyMBEL891 ^19^.

### Metabolic modelling

The reconstructed model was analysed using COBRApy-0.4.1 COnstraints-Based Reconstruction and Analysis for Python ^33^ and COBRAToolbox-2.0 ^34^ in MATLAB^®^ R2016a (Mathworks Inc.). Gurobi 6.5.0 (http://www.gurobi.com) and GLPK 4.32 (http://www.gnu.org/software/glpk/) solvers were used for computational simulations presented. A MATLAB^®^ script to obtain all the results shown in this manuscript is provided as Supplementary Material S8.

Two growth media were considered during the *in silico* simulations:

#### Rich medium

Lower bounds of salts uptake reactions were set to −1000 mmol/g*h^−1^ in order to mimic non-limiting conditions. Carbon sources uptake reactions were set to −100 mmol/g*h^−1^.

#### Simmons medium

^35^: Lower bounds of all uptake reactions accounting for the nutrients present in Simmons medium (see Supplementary Material S1, Supplementary Table 2), were set to −1000 mmol/g*h^−1^, to mimic non limiting conditions, only the C-source (citrate) was set to −5.

### FVA

FVA analysis allows the determination of the span of possible flux variability (*i.e*. the maximum and minimum values of all the fluxes that satisfy the given constraints) while keeping the same optimal objective value.

This approach has been used in this work in order to impose bounds to FBA flux predictions, which are notably non-unique. In fact, for any optimal solution found through FBA there may exist alternate flux distribution patterns yielding the same growth rate. Hence, the space of reliable FBA-flux predictions has been restricted by selecting only those that occur in the interval defined as follows:

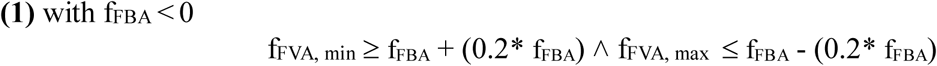

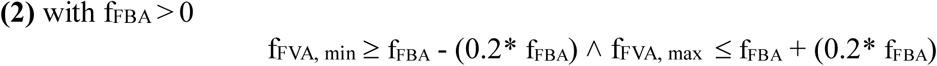

### Gene essentiality and flux ratios calculation

Gene essentiality testing was performed by simulating deletion of each gene within the metabolic network and hence setting the associated reactions to carrying no flux (according to the corresponding Gene-Protein-Reaction (GPR) rule). To predict the growth of the mutant strain and determine the set of EGs, we used two different approaches, FBA and MOMA ^36^. The main difference between them is that while the first predicts growth yield and metabolic fluxes based on the biological assumption of optimal growth, the second does not assume optimality of growth but approximates metabolic phenotype by performing distance minimization in flux space. The second approach has been shown to be more accurate in predicting lethal phenotypes ^36^. The knocked-out gene was defined as ‘essential’ according to the results obtained computing the ratio (GR_ratio_) between the simulated knocked out strain growth rate (*μ*KO) and the one predicted for the wild type strain (*μ*WT). Formulated as:

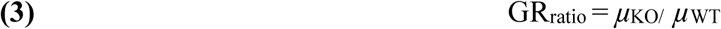

Following this approach, if GR_ratio_ = 0, then the knocked out gene is labelled as essential.

Conversely, in case GR_ratio_ = 1, the removal of the gene has no effect on the growth phenotype. Finally, when 0 < GR_ratio_ < 1, the deleted gene was labelled as fitness-contributing gene, *i.e*. its removal partially affects the capability of the cell to produce biomass.

As MOMA and FBA predictions may lead to different essential gene sets ^37, 38, 36^, we used both approaches to compute essential genes in all the conditions tested in this work. Although no major differences were observed, results obtained with both methods are presented throughout the manuscript.

In order to evaluate the range of the change in the carried flux of each reaction in the model following colistin exposure, we compute the ratio between the predicted flux in the treated *vs*. the untreated conditions as follows:

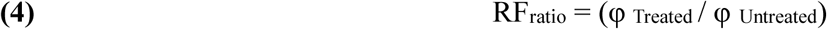

Values of RF_ratio_ equal to one indicate that no changes in the activity of the corresponding reactions were observed when simulating growth in the treated *vs*. the untreated conditions. Conversely, values of RFratio between 0 and 1 or values greater than 1 will indicate a reduced or increased activity of the corresponding reactions in the treated condition, respectively. Finally, negative values of RF_ratio_ will indicate those reactions whose directionality is predicted to change after the treatment.

### Transcriptomics data integration and data visualization

In order to add transcriptional regulatory rules to the metabolic model, we imported the model from COBRA Toolbox into TIGER-1.2.0 0 (Toolbox for Integrating Genome-scale metabolism, Expression, and Regulation) framework (12). Then, the up- and down-regulation ratios of gene expression were mapped into the *A. baumannii* ATCC 19606 metabolic model by using MADE (Metabolic Adjustment by Differential Expression)^24^. The program uses significant statistical changes in gene or gene expression to create functional metabolic models. By adopting an optimization approach that applies Boolean rules, MADE connects reactions to the binary expression states of associated genes. The four arrays of genes to be switched-off yielded by MADE have been reported in Supplementary Material S9.

### EGs BLAST searches in *H. sapiens* and *A. baumannii* species

Protein sequences of the corresponding EGs found in *A. baumannii* ATCC 19606 were probed against the human proteome to test their validity as potential drug target in infections with this pathogen, *i.e*. to exclude any cross-interactions between the drug used for the treatment and human proteome elements.

Queries were aligned to the protein sequences of *H. sapiens* using the default search parameters of the NCBI BLASTP online tool (BLOSUM62 matrix and gap costs equal to Existence 11, Extension 1). Results were considered positive (orthologous sequences found) if their sequence identity score value was equal to/greater than 30.

In addition, the global distribution of EGs was evaluated at the *A. baumannii* species level by probing them against all of sequenced genomes retrieved at NCBI ftp site, *i.e*. 1099 genomes. Particularly, the focus was centred on EGs found in Simmons medium and in rich medium, as well as for those found after 15 and 60 minutes of colistin exposure. BLAST search parameters and analysis of the results were performed as described above.

## Author contribution statement

MF and LP conceived the study and prepared the first draft of the manuscript. LP performed metabolic network reconstruction. LP and MF performed the simulations with the model. LM performed model gap-filling and participated in the modelling step. MF, LP, RF, LD and EB discussed the results and participated in the writing process.

